# Transcranial cortex-wide imaging of murine ischemic perfusion with large-field multifocal illumination fluorescence microscopy

**DOI:** 10.1101/2023.11.01.564959

**Authors:** Zhenyue Chen, Quanyu Zhou, Jeanne Droux, Yu-Hang Liu, Chaim Glück, Irmak Gezginer, Matthias Wyss, Hikari AI Yoshihara, Diana Rita Kindler, Bruno Weber, Susanne Wegener, Mohamad El Amki, Daniel Razansky

## Abstract

Ischemic stroke is a common cause of death worldwide and a main cause of morbidity. Presently, computed tomography and magnetic resonance imaging are the mainstay for stroke diagnosis and therapeutic monitoring. These modalities are often limited in terms of accessibility as well as their ability to map brain perfusion with sufficient spatial and temporal resolution, particularly in the context of preclinical studies, thus calling for development of new brain perfusion techniques featuring rapid imaging speed, cost-effectiveness, and ease of use. Herein, we report on cortex-wide perfusion imaging in murine ischemic stroke with large-field multi-focal illumination fluorescence microscopy (LMI). We attained quantitative readings of hemodynamic and structural changes in cerebral vascular network and pial vessels at capillary level resolution and 80 Hz frame rate fully transcranially. The *in vivo* perfusion maps accurately delineated the ischemic core and penumbra, further exhibiting strong correlation with *ex vivo* triphenyl tetrazolium chloride staining. Interestingly, monitoring of therapeutic effects of thrombolysis in stroke has revealed that early recanalization could effectively save the penumbra whilst reducing the infarct area. Furthermore, cross-strain comparison of perfusion dynamics affirmed that C57BL/6 mice, benefiting from more extensive pial collateral recruitment, feature a larger penumbra and smaller infarct core as compared to BALB/c mice which have few or no collaterals. Finally, we apply LMI to show that sensory stimulation-based treatment enhances blood flow and abolish perfusion deficit in the ischemic core and penumbra regions. The simple, cost-effective and minimally invasive nature of the proposed approach offers new venues for brain perfusion research under various disease conditions such as stroke, neurodegeneration or epileptic seizures.

## Introduction

Acute ischemic stroke is the second cause of death and first cause of morbidity worldwide ^[1]^. Emerging evidence indicates that best outcomes are achieved when recanalization therapy is performed as soon as possible after the stroke onset. During this short period, known as the “Golden Hour”, rapid determination of both reversibly (penumbra) and irreversibly (core) injured tissue is paramount ^[2-5]^. Without early recanalization, the penumbra becomes infarcted whilst late recanalization may increase the rate of intracerebral bleedings and mortality ^[6]^. Hence, fast identification of the ischemic penumbra and its expansion over time is crucial for assessing treatment benefits for stroke patients. However, precision techniques for imaging the penumbra and brain perfusion alterations are still lacking.

X-Ray computed tomography (CT) and magnetic resonance imaging (MRI) are the mainstay for stroke diagnosis and therapeutic monitoring in both preclinical and clinical settings. CT has a proven track record for assessing acute ischemic stroke and ruling out hemorrhage, yet it uses ionizing radiation and suffers from inferior sensitivity and soft tissue contrast ^[1]^. While MR angiography (MRA), diffusion- (DWI) and perfusion-weighted (PWI) imaging can provide comprehensive multiparametric assessment of the brain parenchyma and circulation ^[7]^, those diagnostic procedures are not widely and immediately accessible. Furthermore, when it comes to small animal studies, preclinical imaging modalities are generally limited in their ability to map brain perfusion with sufficient spatial and temporal resolution.

Optical imaging techniques have been explored as an accessible and cost-effective alternative for stroke studies in rodents. Although *ex vivo* histological imaging of brain sections allows for detailed evaluation of stroke injury, including neuronal apoptosis, necrosis and death, this technique lacks dynamic temporal information, requiring post-mortem indirect inferences ^[8]^. Conventional *in vivo* widefield fluorescence imaging suffers from poor spatial resolution due to strong light scattering in the skull and brain tissue, making the assessment of perfusion and single vessel circulation unattainable. On the other hand, point scanning microscopy techniques, such as laser scanning confocal microscopy, optical coherence tomography (OCT) or multi-photon microscopy (MPM), can achieve superior spatial resolution but commonly involve highly invasive craniotomy procedures ^[9, 10]^. Those techniques further suffer from restricted field of view (FOV), making them unsuitable for monitoring the vascular structural change on a brain-wide scale. There is an increasing number of studies exploring the potential of near-infrared spectroscopy (NIRS) due to its relatively deep penetration and non-invasiveness ^[11]^. Although NIRS can partially map superficial cerebrovascular responses, its value for acute stroke imaging is greatly limited by poor spatial resolution and low specificity. Other optical imaging techniques such as time-resolved absolute quantification methods^[12]^, ICG-based perfusion assessment^[13]^ and laser speckle contrast imaging (LSCI) ^[14]^ have been exploited in intra-operative applications. However, these techniques provide poor spatial resolution due to strong scattering or otherwise limited functional information in terms of blood flow and perfusion constant for assessing acute ischemic stroke.

In this work, we introduce large-field multi-focal illumination (LMI) fluorescence microscopy for transcranial cortex-wide imaging of microvasculature, functional dynamics, and perfusion maps in ischemic stroke with high spatial and temporal resolution. *In vivo* assessment of ischemic core and penumbra and monitoring of recanalization therapies is performed with the method, depicting the process of rescuing the penumbra via reperfusion by thrombolysis. The *in vivo* data is validated with corresponding triphenyl tetrazolium chloride (TTC) staining of brain infarctions. Furthermore, we performed cross-strain comparison of post-stroke perfusion dynamics in mice with different vascular architectures (C57BL/6 and BALB/c) and revealed different core and penumbra sizes as well as collateral recruitment. Finally, sensory stimulation-based treatment for stroke, as a proof of concept, is performed and monitored with the proposed method to check its therapeutic effect.

## Results

### *In vivo* transcranial fluorescence microscopy allows noninvasive brain-wide perfusion imaging in ischemic stroke

The LMI technique capitalizes on the fast scanning based on an acousto-optic deflector (AOD) and a beam-splitting grating to generate a uniform illumination grid of 17×17 foci (Fig. 1**a**). By rapidly scanning the illumination grid and synchronizing data acquisition with a high-speed camera, a combination of a large FOV of 7×7 mm^2^ and high spatial resolution of 14.4 µm and high frame rate of 80 Hz was achieved, thus exceeding imaging performance as compared to previously reported implementation ^[15, 16]^. The complete experimental procedure included several steps. First, ischemic stroke is induced by injecting thrombin into the right middle cerebral artery (MCA) as described previously ^[17, 18]^. Subsequently, time-lapse LMI images are acquired and compared with the images recorded with the well-established LSCI technique. For the LMI imaging, Sulfo-Cyanin-5.5-carboxylic-acid (Cy5.5) dye is administrated intravenously through the tail vein. For treated animals with thrombolysis, t-PA (0.9 mg/kg) was administered as described previously ^[19]^after LMI imaging, i.e., 60 min post-stroke. For sensory stimulation treatment, hindpaw electrical stimulation was performed (see Methods).

**Figure 1.**
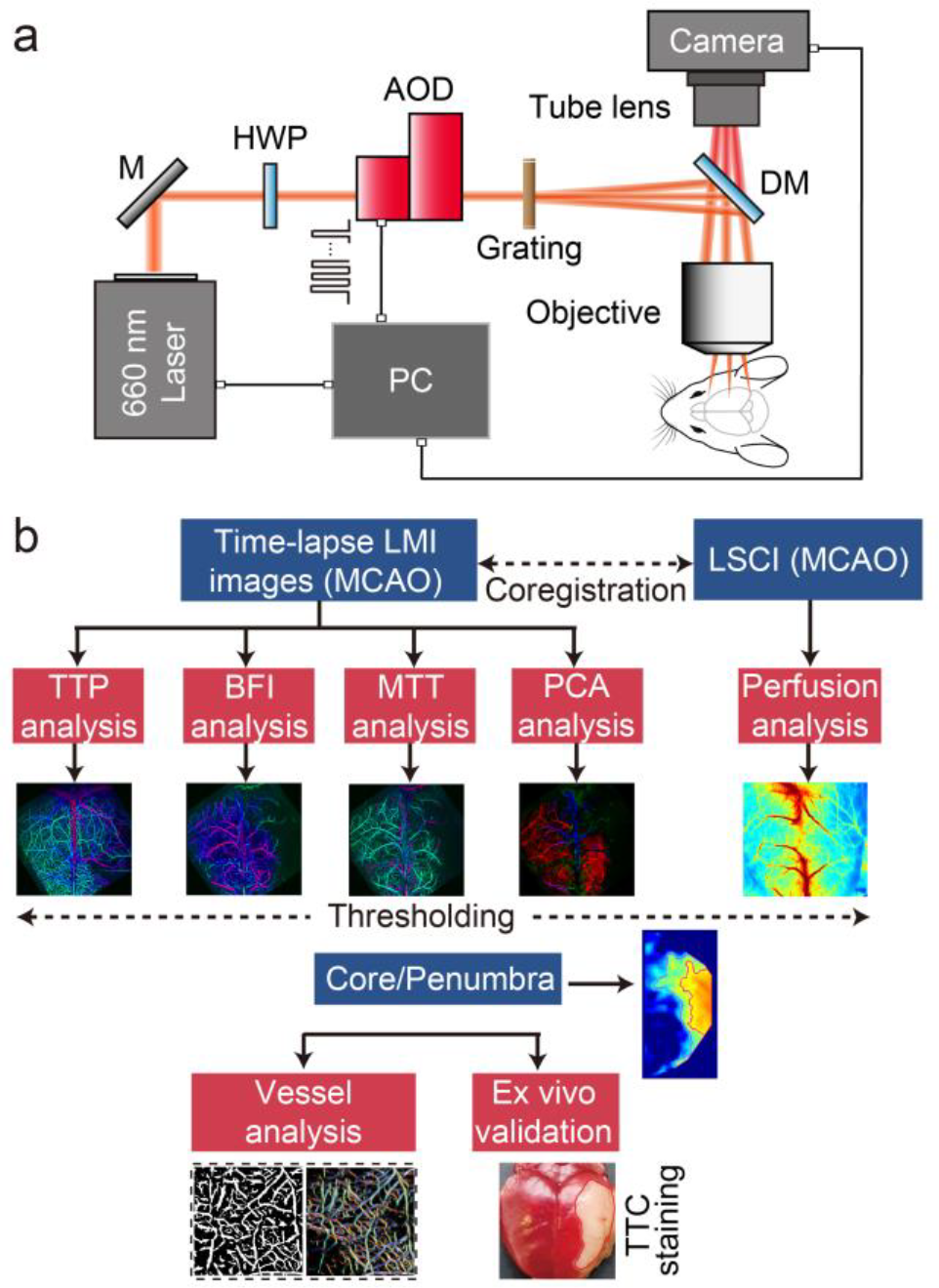
Stroke imaging system and methodology. **a.** Layout of the large-field multi-focal illumination (LMI) fluorescence microscopy system. **b**. Data processing pipeline of LMI and comparison to conventional laser speckle contrast imaging (LSCI) method.

An offline data processing pipeline was developed to facilitate stroke assessment (Fig. 1**b**). LMI images were first reconstructed and subsequently co-registered with the LSCI images. Pixel-wise time to peak (TTP), blood flow index (BFI) and mean transient time (MTT) maps were calculated from the LMI image stack. Principle component analysis (PCA) was performed to the image stack to distinguish the skull vessels and brain vessels (arterial and venous vessels) based on their different perfusion profiles. Blood flow velocity and cerebral blood volume were calculated based on regions of interest (ROIs) in different vessels. For more details on the calculation, please check the Methods section and Fig. S1. Infarct core and penumbra regions were obtained after applying thresholds to the BFI signal changes by comparing the ipsilateral and contralateral hemispheres. Note that the cortical blood flow from the contralateral hemisphere remained unchanged from the baseline following the stroke surgery, as verified with fluorescence localization microscopy (Fig. S2), which is also in line with previous reports ^[20]^. ROIs including core and penumbra were selected to perform further statistical analysis on perfusion constants and vessel fill fraction. Finally, validation of the *in vivo* measurement was performed by TTC staining which measures ischemic infarction in the isolated brains.

Mouse brains were imaged (n = 3 C57BL/6 mice) with LMI microscopy pre- and post-stroke onset to map the cerebral microcirculation (Supplementary videos 1, 2). PCA results prior to the stroke induction corroborated anatomical information (Fig. 2**a**) on the cerebral vascular network and revealed clear perfusion differences between arteries and veins (Fig. 2**e**). Post-stroke PCA corroborated the occlusion of MCA while other vessels such as anterior cerebral arteries (ACAs) were not affected (Fig. 2**k**). BFI, MTT and TTP maps in a representative dataset showed clear perfusion deficit after stroke (Figs. 2**b-d, h-j**). These results were further validated by independent measurement with LSCI which showed similar patterns of perfusion deficits (Figs. 2**f, l**).

**Figure 2.**
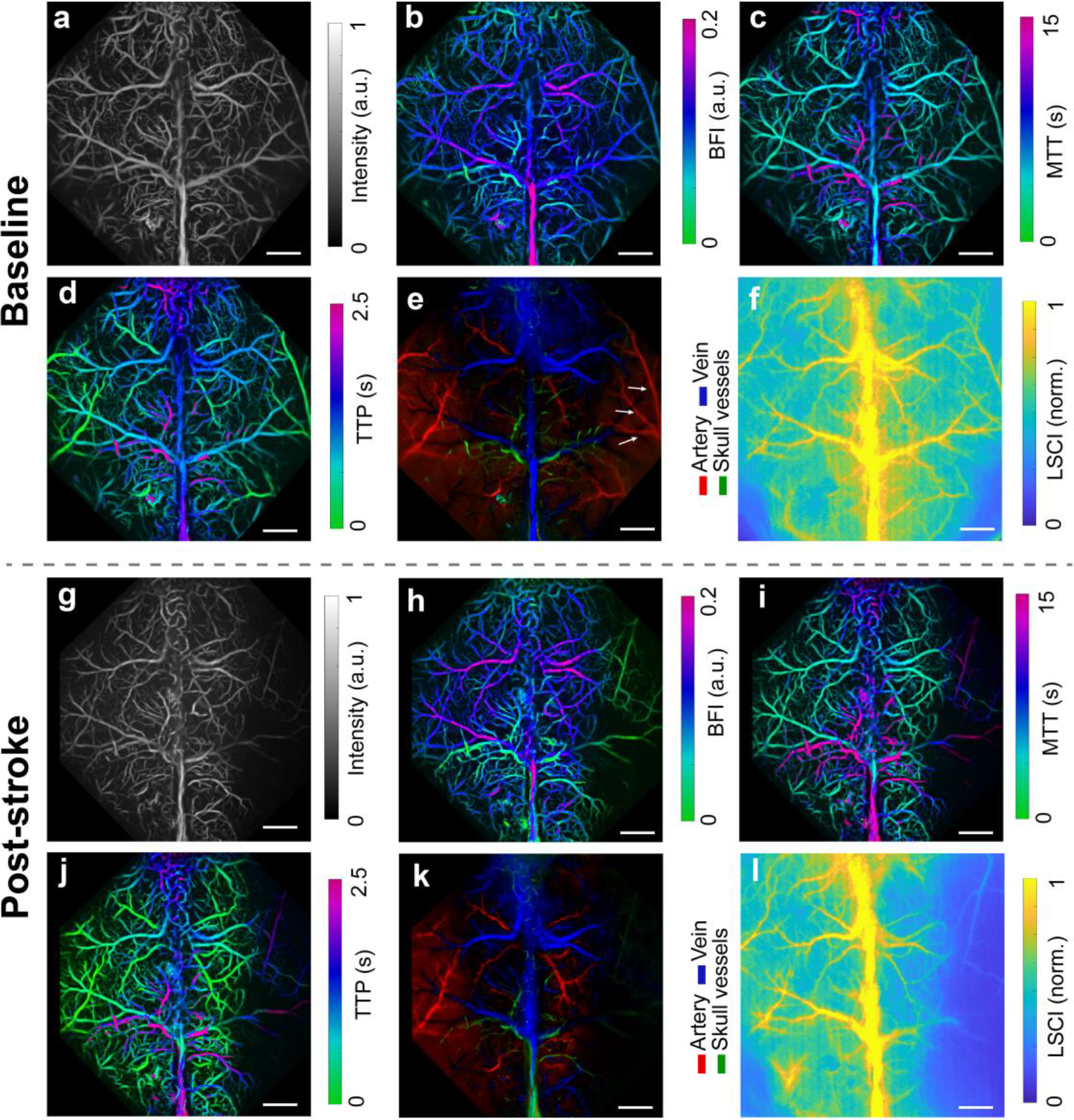
Transcranial LMI fluorescence microscopy imaging allows detection of perfusion deficits in mouse stroke models. **a** – **e** Anatomical image, BFI, MTT, TTP and PCA results from a representative mouse brain prior to stroke. **f** The corresponding LSCI image. **g** - **l** Respective images acquired from the same animal after ischemic stroke. All scale bars: 1 mm. The experiment was repeated independently in 3 mice with similar results.

### LMI fluorescence microscopy allows the prediction of therapy success in stroke

After the stroke induction, ischemic lesion boundaries detection, including discrimination between the core of lesion and penumbra, could be accomplished with LMI (Figs. 3**a, b**). To demonstrate the precision of our proposed method, we further assessed the effect of a thrombolytic drug tPA, which allows CBF recovery and penumbra salvage. Mice were categorized into two different groups: stroke + tPA versus stroke controls (n=3 BALB/c mice per group). Before treatment, fluorescence intensity, BFI as well as fill fraction did not differ significantly between the two groups of mice. After tPA infusion, statistics on fluorescence intensity (Fig. 3**c**), BFI (Fig. 3**d**), LSCI (Fig. 3**e**), lesion size estimation (Fig. 3**f**) and vessel fill fraction (Fig. 3**g**) clearly show improved reperfusion both in the core and penumbra regions post-treatment. Interestingly, in the mice treated with tPA, TTC staining revealed the infarction area was highly correlated with the core region detected by LMI, meaning the penumbra was saved (Fig. 3**b**). Furthermore, in the control mice, TTC staining revealed a large infarct combining both core and penumbra detected by LMI (Fig. 3**a**). Compared to LSCI, correlation coefficients between the in vivo assessed core region and the TTC stained infarct tissue reveal that LMI has significantly higher diagnostic accuracy (86.1 ± 4.5% versus 75.4 ± 5.2%, MEAN ± STD, p = 0.0032) (Fig. 3**h**). These data confirm the precision of LMI microscopy in discriminating penumbra from the core.

**Figure 3.**
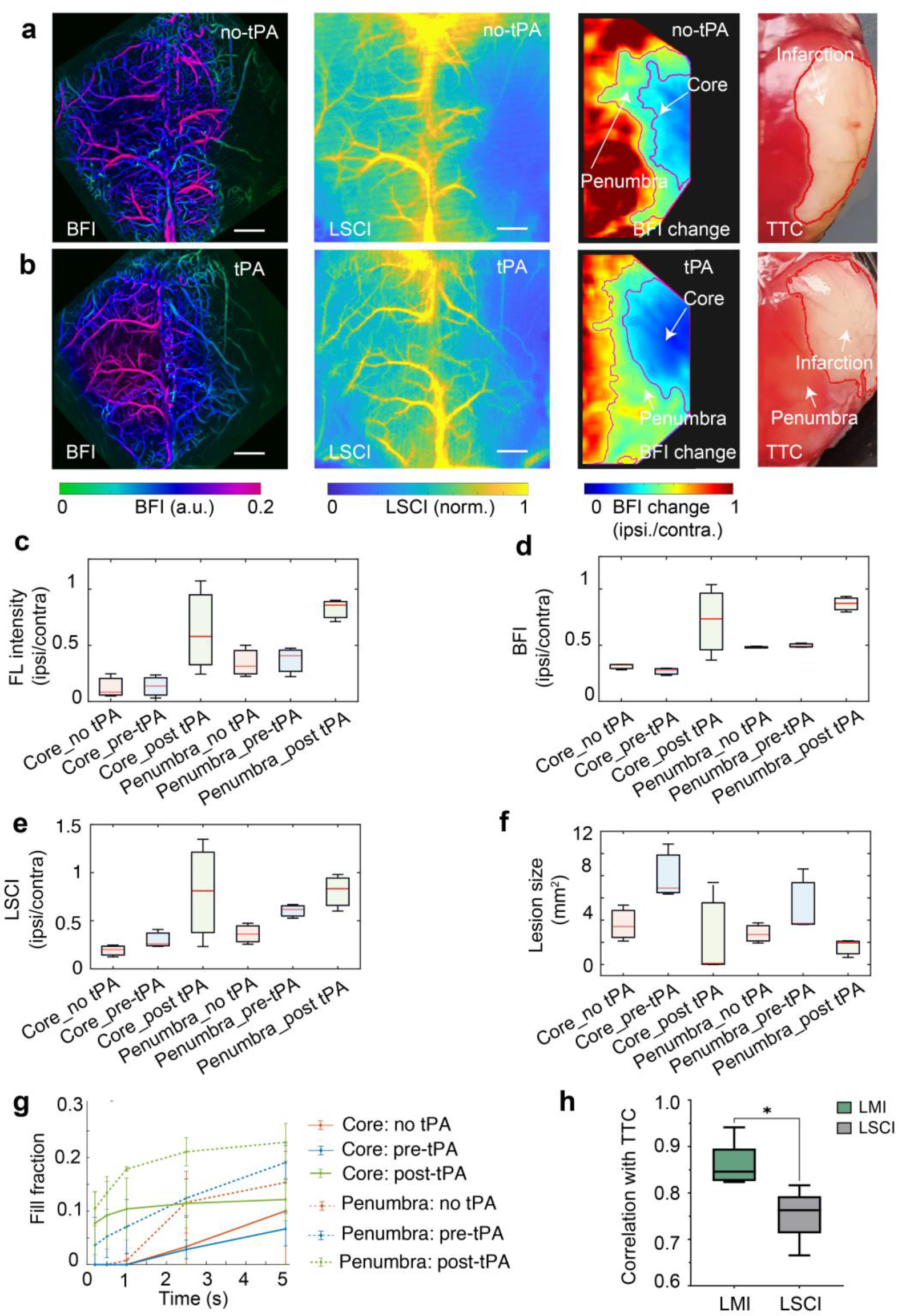
Stroke imaging results w/o tPA treatment in BALB/c mice. **a, b** BFI, LSCI, BFI change and corresponding TTC staining from the representative mouse without and with tPA treatment, respectively. **c**-**f** Comparison between the LMI fluorescence intensity, BFI, LSCI and lesion size for non-treated, pre-treatment and post-treatment mice. **g** Vessel fill fraction comparison between nontreated, pre-treatment and post-treatment mice. **h** Infarct size correlation between *in vivo* measurement with LMI/LSCI and TTC staining. Statistics performed with paired two-tailed t-test. All scale bars: 1 mm. The experiment was repeated independently in 3 mice per group with similar results.

### LMI fluorescence microscopy reveals strain-related perfusion patterns post-stroke

Individual susceptibility to ischemic injury as well as the size of the ischemic penumbra vary greatly among different strains of mice due to the differences in their number of leptomeningeal collaterals. In this experiment, we aimed to perform LMI in animals with different core/penumbra ratio. To do so, we induced stroke in mice with different collateral networks, namely, BALB/c having poor collaterals and C57BL/6 having rich collaterals (n = 3 in each group). BFI, MTT and TTP maps show clear difference between these two strains (Figs. 4**a** vs 4**b** and Figs. S3). Specifically, BALB/c mice were more susceptible to the ischemic injury induced by stroke than C57BL/6 mice. Furthermore, BALB/c mice showed significantly reduced blood circulation in the infarct region (Fig. 4**a**) with partially reduced blood flow (Fig. 4**b**) and higher time latency (Figs. 4**e**) observed in the infarct region of C57BL/6 mice. Such discrepancy is congruent with previous reports and can be attributed to the lack of collateral vessels in BALB/c mice ^[21]^.

**Figure 4.**
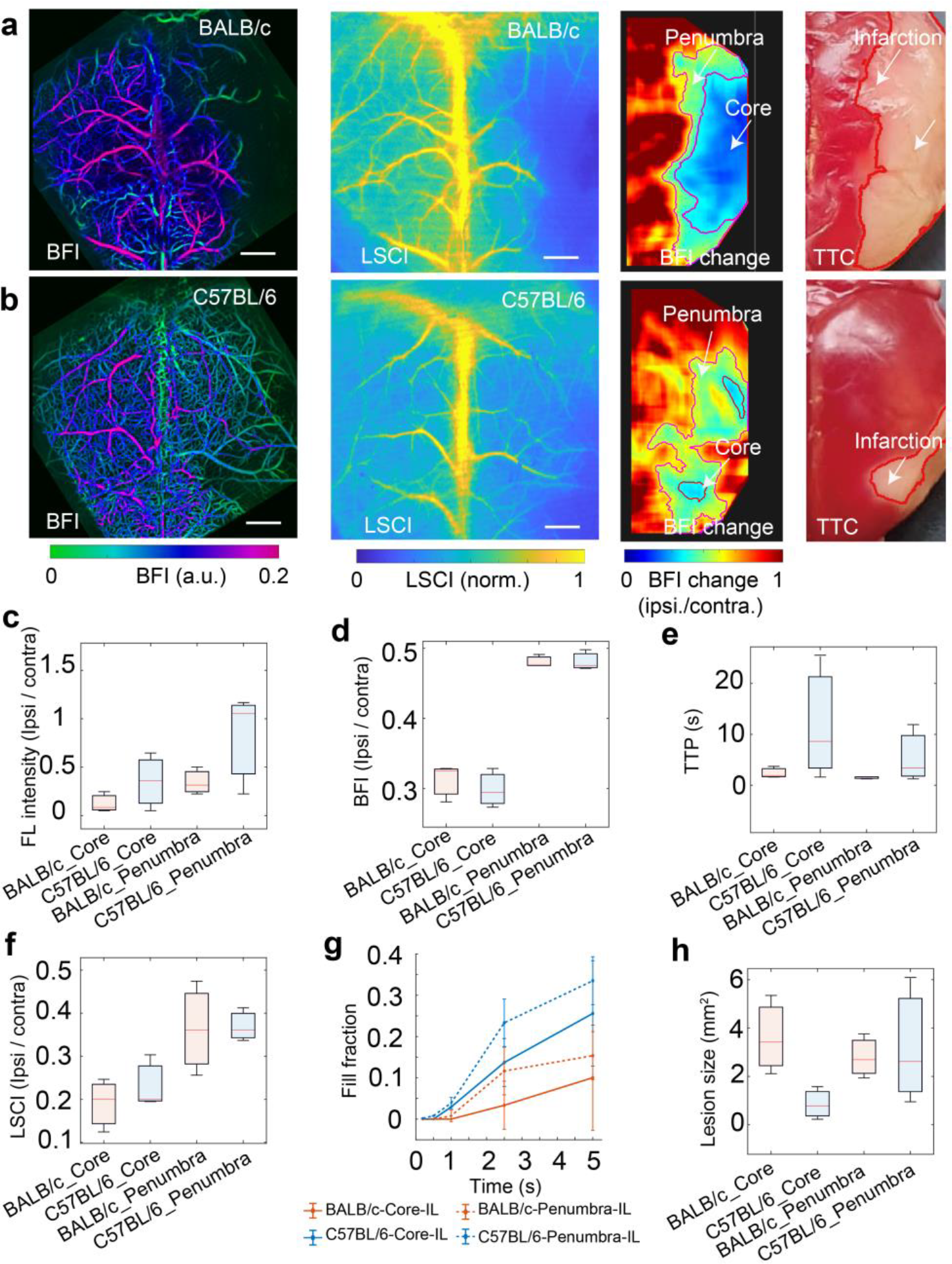
Experimental results on stroke imaging in BALB/c and C57BL/6 mice. **a, b** BFI and LSCI images along with BFI signal change and TTC stainings from representative BALB/c and C57BL/6 mice. **c-f** Statistics on fluorescence intensity, BFI, TTP, LSCI from both BALB/c and C57BL/6 mice. All the parameters were normalized to values from the contralateral side. **g** Vessel fill fraction time course in the core and penumbra regions from both BALB/c and C57BL/6 mice. **h** Statistics on the lesion size. All scale bars: 1 mm. The experiment was repeated independently in 3 mice per group with similar results.

The above observations are generally corroborated by the LSCI results (Figs. 4**a, b**, 2^nd^ column). However, in comparison to LSCI, the core and penumbra regions inferred from the LMI measurements (Figs. 4**a** and 4**b**, 3^rd^ column) manifested significantly higher correlation with the TTC stained brain tissue (Figs. 4**a** and 4**b**, 4^th^ column, and Fig. S4). This is arguably ascribed to the unspecific signal contrast mechanism of LSCI which chiefly represents a non-linear dependence between tissue scattering and blood flow velocity ^[22]^. Note that TTP and MTT maps also manifest a clear boundary between the healthy and infarcted tissue. This is of importance since the infarcted areas have prolonged TTP and MTT values due to impaired microcirculation, which can serve as additional criteria for stroke assessment without requiring a priori information about blood circulation or comparison with the other brain hemisphere. Yet, TTP and MTT maps are inferior in distinguishing the infarct core and penumbra (Figs. S5 and S6). Quantitative assessment of perfusion alterations was further performed for both LSCI and LMI images in the infarct core and penumbra regions. Statistics on fluorescence signal intensity (Fig. 4**c**), BFI (Fig. 4**d**), and LSCI (Fig. 4**f**) show that BALB/c mice exhibited reduced perfusion post-stroke as compared to C57BL/6 mice whilst C57BL/6 mice had longer perfusion duration (Fig. 4**e**). Consequently, BALB/c mice had larger infarct core size than C57BL/6 mice (Fig. 4**h**). Structural information analysis on vessel fill fraction also reveals that C57BL/6 mice had higher vessel perfusion rate at all sampling time instants as compared to BALB/c mice, which is consistent with aforementioned observations (Fig. 4**g**).

### LMI fluorescence microscopy captures dynamic collateral recruitment

To understand the impact of collaterals in maintaining brain perfusion after stroke, we compared the collateral circulation between C57BL/6 mice and BALB/c mice. Time-lapse images from a selected ROI covering both ACA and MCA were recorded from one representative mouse of each strain. Under baseline healthy conditions, collateral recruitment was not observed suggesting that collateral flow was low (Fig. 5**a, b**, 1^st^ row). Following stroke induction, collateral flow became pronounced in C57BL/6 mice (Fig. 5**a**, 2^nd^ row) but remained unapparent in BALB/c mice (Fig. 5**b**, 2^nd^ row). Note that all the experimental parameters were kept the same for the assessment of both strains.

**Figure 5.**
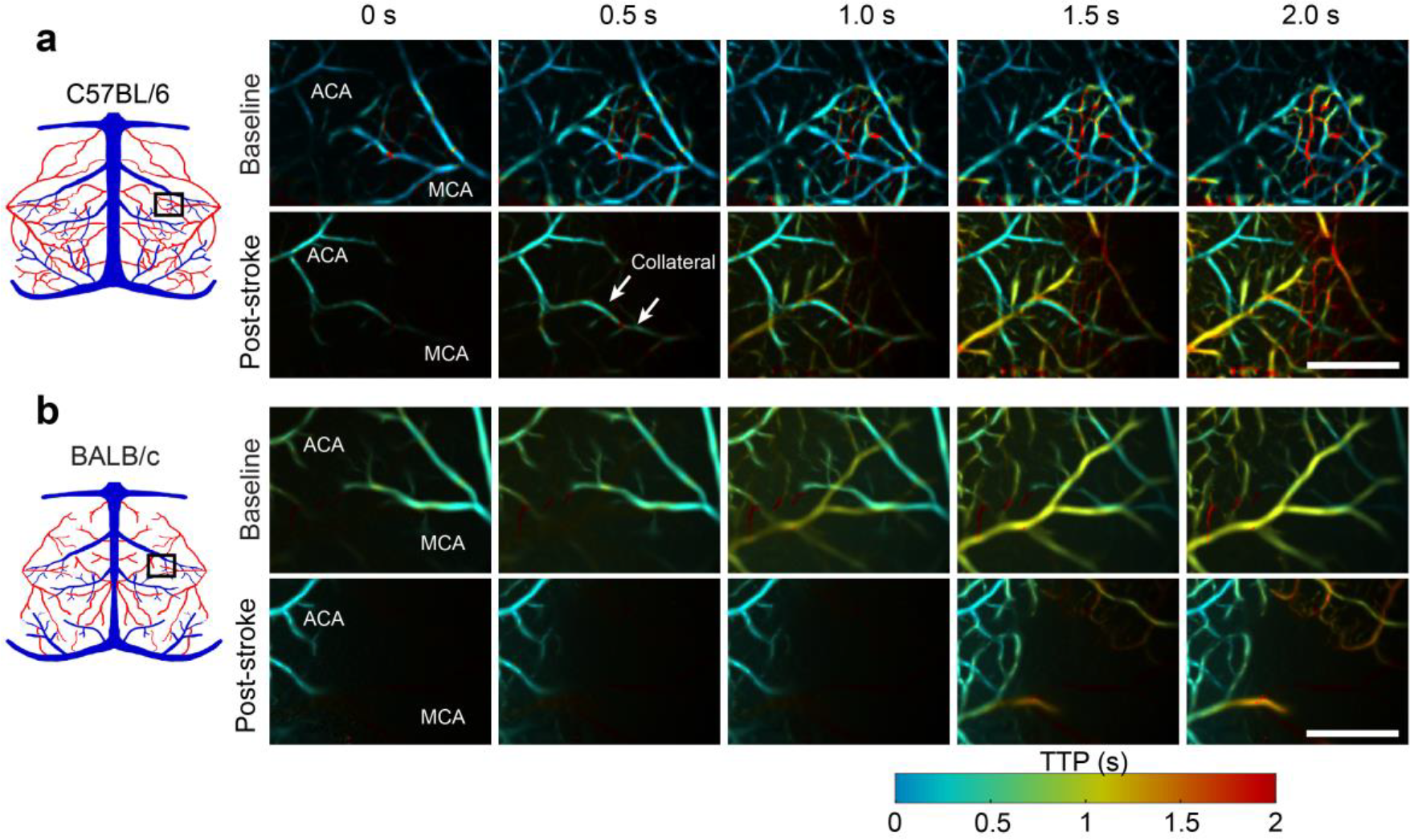
Dissimilar perfusion dynamics in C57BL/6 mice and BALB/c mice. **a** Representative timelapse images post fluorescence dye injection into C57BL/6 mouse. Top row: baseline before stroke; bottom row: after stroke; diagram on the left illustrates the imaged ROI. **b** Representative time-lapse images post fluorescence dye injection into BALB/c mouse. Top row: baseline before stroke; bottom row: after stroke; diagram on the left illustrates the imaged ROI. All scale bars: 1 mm. The experiment was repeated independently in 3 mice per group with similar results.

### Sensory stimulations, delivered within one hour of the stroke onset, protect the cortex from perfusion deficits

We finally investigated whether a sensory stimulation-based treatment can be exploited to improve cortical reperfusion after stroke. To this end, a sensory stimulation trial was performed immediately after the stroke onset (Fig. 6**a**). The same experimental procedure was followed for animals from both the intervention (n = 7) and the control (n = 4) groups (see more details in the Methods section). Prior to inducing stroke, cortical activity and blood flow in the S1HL cortical region were assessed with intrinsic signal optical imaging (IOI) and LSCI, respectively, whilst brain perfusion was measured with LMI. To test the effect of sensory stimulation on post-stroke brain perfusion, LMI and LSCI were repeated immediately after stroke (0h) and at the end of the sensory-stimulation-based treatment (1h post-stroke).

**Figure 6.**
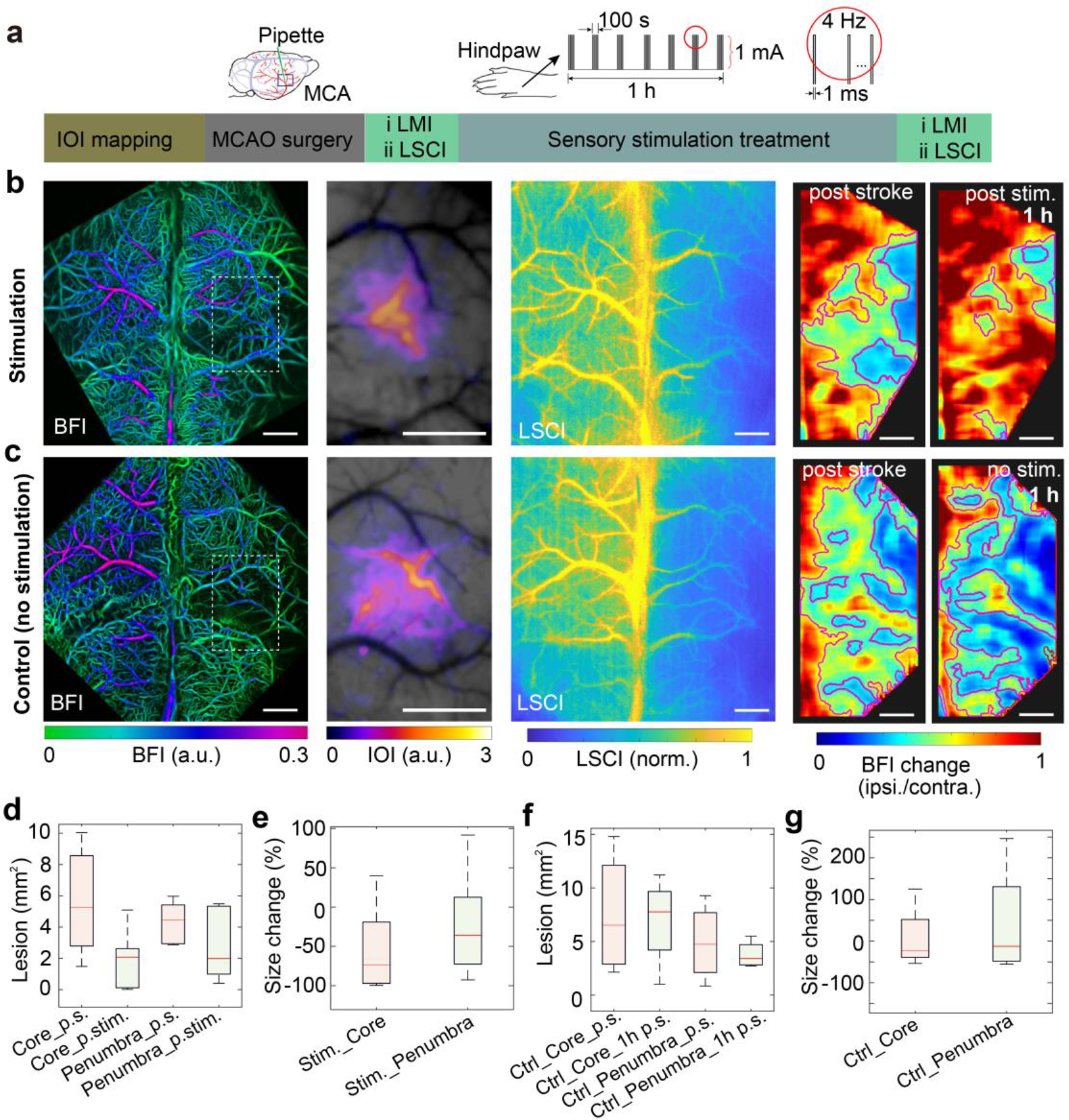
Effects of sensory stimulation treatment post stroke in C57BL/6 mice. **a** The experimental pipeline. **b** Imaging results from one representative mouse with hindpaw stimulation. From left to right: BFI post-stimulation, intrinsic signal (IOI) mapping of the stimulated S1HL region, LSCI poststimulation, BFI change post-stroke and post-stimulation. **c** Corresponding results from a representative control mouse post-stimulation. **d** Core and penumbra size post-stroke and poststimulation. **e** Relative changes of core and penumbra size after hindpaw stimulation. **f** Core and penumbra size 0 h and 1 h post-stroke in control mice. **g** Relative changes of core and penumbra at 1 h post-stroke. All scale bars: 1 mm. The experiment was repeated independently in 7 mice for sensory stimulation treatment and 4 mice for control with similar results.

BFI and LSCI (Fig. 6**b**) displayed decreased hypoperfused area in the stimulated mice; however, the brain perfusion deficit was not different between the 0 h and 1 h post stroke time points in the nonstimulated mice (Fig. 6**c**). Note that the perception of the lesion size from BFI map may be affected by the display of the image after contrast adjustment. Instead, ameliorating brain perfusion was evident when evaluating the BFI changes (Fig. 6**b**). In contrast, the perfusion deficit has intensified at 1 h poststroke for control mice by evaluating the BFI changes (Fig. 6**c**). Statistics on the lesion size and its relative deviation from the intervention group show a clear contraction of the core region (p = 0.0247) and a tendency for size reduction in the penumbra (p = 0.2295) whilst the lesion sizes of both core and penumbra remained nearly unaltered in the control group (Figs. 6**f, g**).

## Discussion

In ischemic stroke, both vascular morphology as well as blood flow information are crucial for accurately discerning the infarct core and penumbra. LMI imaging brings major advantages for the detection and quantification of the structural and functional alterations after stroke, as well as for precise assessment of the core and penumbra regions. The method attains a powerful combination between high (capillary-level) spatial resolution, millisecond temporal resolution, transcranial imaging capacity and whole-cortex coverage. It is observed that blood flow in the infarct core is impaired to a greater extent as compared to the penumbra. Interestingly, the technique has been used to predict the success of thrombolytic therapy and a sensory stimulation-based treatment. Furthermore, our study shows how LMI can be used to quantify differences in brain perfusion dynamics between mice with different stroke severities. Collateral recruitment, serving as an alternative blood supply to the infarct region, is of great interest in stroke as it crucially determines infarct severity, treatment efficacy and recovery. In this work, by real-time monitoring of cortex-wide microcirculation, collateral recruitment was observed in C57BL/6 mice post-stroke while no similar phenomenon was seen in the baseline (pre-stroke) measurements or in BALB/c mice.

In preclinical stroke research, LSCI has been widely adopted to provide brain-wide microcirculation mapping in real-time and can indicate abnormal blood flow in different brain regions. However, the spatial resolution of LSCI is insufficient for accurately determining alterations in specific vessels. Furthermore, the overall ability of LSCI to provide quantitative readings of absolute blood flow, velocity, or tissue perfusion remains questionable ^[23]^. Interestingly, LMI provides superior performance by achieving higher spatial and temporal resolution as evinced by high resolution mapping of the cerebral vascular network and corresponding perfusion dynamics recorded at a frame rate of 80 Hz. More importantly, unlike LSCI whose imaging contrast mainly stems from scattering and is indirectly related to blood flow, the LMI approach can image the blood flow directly thus ensuring a higher interpretation power of the results and providing a higher correlation to the TTC staining which further correlates well with immunohistochemistry using NeuN and DAPI staining (Fig. S7). BFI, TTP and MTT values obtained with the proposed LMI approach were further validated with dynamic contrast-enhanced perfusion MR imaging using superparamagnetic iron dioxide nanoparticles (SPION) injected through the tail vein (Figs. S8 and S9). By applying the same data analysis to ROIs in different types of vessels, i.e., MCA, drainage veins, superior sagittal sinus (SSS) and rostral rhinal vein (RRV), no significant difference was found in the BFI/TTP/MTT values from each vessel type between LMI and MRI, except BFI and MTT from SSS, where fluorescence data presented lower BFI and prolonged MTT arguably caused by the fluorescence signal integration across different depths in the cortex and skull (Fig. S9).

To this end, CT and MRA have successfully been employed both in pre-clinical and clinical stroke studies. However, X-Ray CT angiography suffers from low spatio-temporal resolution and has been shown to result in overestimation of the infarct size ^[1, 24]^. CT perfusion studies can produce high spatial resolution quantitative mapping of cerebral blood volume, flow, and TTP at the expense of longer acquisition time, radiation dose and contrast agent dose ^[1]^. MRA can identify the location and extent of occlusive thrombus and identify tandem lesions at the cervical level ^[1]^. Yet, its outcomes are often compromised by the prolonged scanning times and other well-known safety issues exacerbated in the acutely unwell stroke patients. Although NIR fluorescence imaging suffers from limited penetration as compared to CT and MRI, it has been adopted for various surgical interventions, including vascular, gastrointestinal, cardiac, and reconstructive surgery ^[25-27]^. The proposed method could potentially improve the quality of NIR fluorescence imaging and gain additional information for better perfusion assessment in stroke studies.

The ischemic penumbra has been in the center of stroke research for the last 40 years. The success for stroke therapy, including mechanical thrombectomy and/or intravenous thrombolysis, is based on a careful selection of patients that had salvageable penumbra. Therefore, stroke assessment studies might greatly benefit from a rapid and accurate tool for quantification of the ischemic penumbra. Despite major improvements in imaging methodology and instrumentation, it is widely accepted that the discrimination between penumbra and core regions is challenging with the existing imaging approaches. This is chiefly due to the considerable variability of CBF thresholds used for delineating these regions. For instance, the classical penumbra has been defined as a region of 20-40% of residual CBF ^[28]^. However, other studies defined the penumbra as a region with 30-50% CBF decrease ^[29, 30]^ whilst some studies even considered regions with blood flow up to 60-70% as the penumbra region ^[31, 32]^. In this work, BFI residuals of <40 % and 40-60 % were considered for delineating the core and penumbra regions, respectively.

As an alternative to this hemodynamic concept, additional studies defined the penumbra as the tissue doomed to infarction in the absence of early reperfusion. Our data confirm that the penumbra detected with LMI can be saved from tissue necrosis with reperfusion, otherwise it turns necrotic over time and merges into the infarct core. Several clinical trials have demonstrated the clinical benefits of salvaging the penumbra in stroke patients ^[33, 34]^, underscoring the importance of imaging tools capable of predicting patient’s response to thrombolytic treatments. It has also been shown that patients with a favorable imaging-based profile have strong collaterals and moderate infarct progression ^[35]^. The size of core and penumbra is underpinned by the collateral supply to the tissue. In line with these clinical data, we have shown that LMI can accurately predict the differences between infarcted areas in animals with rich and poor collaterals within the first minute after occlusion. Importantly, since our imaging system allows for transcranial imaging across the entire cortex, it was possible to perform multiparametric characterization of the pial vessels including perfusion dynamics and collaterals recruitment. Clinically, this information will aid in monitoring the role of pial collaterals in infarct development and recovery. Note that the effective imaging depth of LMI is limited by the ability to maintain a focused beam pattern as it propagates through turbid brain tissues, which typically corresponds to 0.5–1 mm usable depth range depending on the wavelength selection. The penetration depth can potentially be improved by performing imaging in the second near-infrared spectral window (NIR-II window, 1000-1700 nm), which benefits from reduced scattering, absorption, and negligible autofluorescence ^[36]^. However, the currently available fluorescent contrast agents in the NIR-II window offer inferior performance in terms of fluorescence quantum-yield and biocompatibility as compared to their counterparts in the visible and NIR-I windows ^[37, 38]^.

Given that evoked neuronal activity from paw stimulation is linked to an increase in blood flow, we further tested whether a potential treatment approach based on sensory stimulation may enhance brain perfusion thus eliminating reperfusion deficits in the core and penumbra regions. It has been shown that sensory stimulation reduces the infarcted region both in the core and penumbra. Enhancing brain reperfusion with sensory stimulation is a particularly promising strategy for stroke treatment because it is non-invasive, nonpharmacological, rapid, and has no side effects. Peripheral sensory stimulation during the hyperacute phase of stroke was reported to significantly recover neural activity ^[39, 40]^. Likewise, sphenopalatine ganglion stimulation has been shown likely to improve functional outcome of acute ischemic stroke patients clinically ^[41]^. Note that when the activated cortex is within a critical range of ischemia, somatosensory activation of peri-infarct cortex may aggravate the situation by increasing the supply-demand mismatch and triggering peri-infarct depolarizations ^[42]^. In the current study, the S1HL region was carefully selected such that it is able to redirect blood flow from ACA to MCA, yet located far enough from the peri-infarct region. The stroke treatments with sensory stimulation were accordingly only performed with mice having rich collateral vascularization.

In summary, we present a high-speed fluorescence microscopy technique for transcranial stroke imaging in mice that allows to (1) confirm ischemic stroke diagnosis, (2) localize the site of vascular occlusion, (3) detect hemodynamic changes and perfusion deficit and (4) identify ischemic core and penumbra with high sensitivity. The method is unique in its ability of localizing brain perfusion changes and accurately assessing the ischemic penumbra. LMI can directly render images of diverse mouse models with local CBF variations. Overall, LMI is simple and cost-effective and may be used to improve our understanding of vascular responses under pathologic conditions, ultimately facilitating clinical diagnosis, monitoring and development of therapeutic interventions for cerebrovascular diseases.

## Methods

### LMI fluorescence microscopy system

The LMI fluorescence imaging method has been adapted from the originally reported system ^[15]^ to provide high spatial resolution and accelerated imaging performance. It is based on a beam-splitting grating and an acousto-optic deflector (AOD) synchronized with a high-speed camera (Fig. 1a). To generate the multifocal illumination, a CW laser at 660 nm (gem 660, Laser Quantum, UK) together with a 17×17 beamsplitting grating (DER243, Holoeye, Germany) were employed. The resulting minibeam pattern was focused by a scan lens (CLS-SL, Thorlabs, USA) to form an illumination grid upon the mouse brain. After passing through the dichroic mirror, the emitted fluorescence signal was collected by an AF micro-Nikkor 60 mm objective (Nikon, Japan) and focused onto the sensor plane of the highspeed CMOS camera (pco.dimax S1, PCO AG, Germany) after passing through a 671 nm longpass filter (F76-671, Semrock, USA) behind the objective. By raster scanning the laser beam with the AOD (DTSXY-400-532, AA Opto-Electronic, France) at a high speed up to hundred kilohertz, high resolution LMI images can be acquired in real-time.

### Animal models

Six BALB/c mice (8 to 9 week-old, Charles Rivers) and fourteen C57BL/6 mice (8 to 9 week-old, Charles Rivers) were used in this work. Animals were housed in individually ventilated, temperature-controlled cages under a 12-hours dark/light cycle. Pelleted food (3437PXL15, CARGILL) and water were provided ad-libitum. All experiments were performed in accordance with the Swiss Federal Act on Animal Protection and were approved by the Cantonal Veterinary Office Zurich (ZH165/2019).

### Thrombin stroke model

The thrombin stroke model was used as described previously ^[17]^. In brief, mice were fixed in a stereotactic frame, the skin between the left eye and ear was incised and the temporal muscle was retracted. A small hole was drilled in the temporal bone to access the MCA while the skull was kept intact. To induce the formation of a clot, 1 μl of purified human alpha-thrombin (1UI; HCT-0020, Haematologic Technologies Inc., USA) was injected into the MCA with a glass pipette. The pipette was removed 10 min after thrombin injection. Ischemia induction was considered stable when cerebral blood flow (CBF) rapidly dropped to at least 50% of baseline level in the MCA territory by checking the LSCI images ^[17-19]^. Note that this threshold was chosen based on our previous experience. We calculated the post-stroke CBF drop for all the mice used in our work and observed 74.2% drop in the mean perfusion values, agreeing well with the conventional thresholds defining stable ischemia induction.

### Laser speckle contrast imaging (LSCI)

Cortical perfusion was monitored post-stroke before and after tPA treatment with a commercial LSCI device (FLPI, Moor Instruments, UK). The LSCI images are generated with arbitrary units in a 16-color palette by the MoorFLPI software.

### Stroke treatment with t-PA administration

t-PA treatment was performed 1 h post-surgery after stroke screening with sequential LMI and LSCI. Human t-PA (0.9 mg/kg, Actilyse, Boehringer Ingelheim) was administrated through tail vein infusion with a syringe pump at a constant infusion speed of 6 µl/min.

### Sensory-stimulation-based treatment for stroke

For stroke treatment with sensory stimulation, 11 C57BL/6 mice were separated into the intervention group (n = 7) and the sham-controlled group (n = 4). All mice were anesthetized with ketamine and xylazine and S1HL cortex region was firstly mapped by using intrinsic optical imaging. Immediately after stroke, perfusion imaging with LSCI and LMI was performed. Mice in the intervention group received electrical hindpaw stimulation for 100 s (pulse duration 1 ms, repetition frequency 4Hz, pulse amplitude 1 mA) with a time interval of 10 min and a duration of 1 hour. Mice in the control group did not receive any stimulation while all other factors (e.g., air supply, heating, and physiological status monitoring) were kept the same as for the stimulated group. During the experiment, an oxygen/air mixture (0.2/0.8 L.min^-1^) was provided through a breathing mask. The body temperature was kept around 37 °C with a feedback-controlled heating pad. Peripheral blood oxygen saturation, heart rate and body temperature were monitored (PhysioSuite, Kent Scientific) in real-time.

### *In vivo* imaging

For *in vivo* imaging, the mice were anesthetized with isoflurane (3.0 % v/v for induction and 1.5 % v/v during experiments) in 20% O_2_ and 80% air at a flow rate of ∼0.5 L/min. The scalp was removed to reduce light scattering. To obtain the cerebral vascular network and the perfusion pattern post-stroke and pre-tPA treatment, each mouse was administrated the Sulfo-Cyanin-5.5-carboxylic-acid (Lumiprobe GmbH, Germany) fluorescent dye solution in PBS (50 µl, 2 mg/ml) through a tail-vein injection while recording with the LMI system, followed by LSCI measurement. Stroke treatment with t-PA administration was performed after the first round of *in vivo* stroke imaging (1 h post-stroke). After finishing the infusion, the mouse was kept in a small animal recovery chamber (Hugo Sachs Elektronik GmbH) for 30 min, followed by the second round of stroke imaging with both LMI and LSCI as described above. For LMI fluorescence microscopy, data were recorded right after fluorescence dye injection and lasted for 20 s. Each LMI image was acquired through 30×30 raster scan with the AOD at ∼11 μm step size, which corresponded to an effective frame rate of 80 Hz. For LSCI, data were recorded for 1 min with an integration time of 250 ms for each LSCI image corresponding to a frame rate of 4 Hz.

### Intrinsic signal optical imaging (IOI)

For intrinsic signal optical imaging, the head of anesthetized mice was shaved and skull exposed after midline incision of overlying skin. The skull was then covered with low melting point agarose in Ringer’s solution (1.5%) and sealed with a glass coverslip. Functional imaging was conducted under 570 nm wavelength illumination (Polychrome V, FEI) using custom-written control software (Matlab version 2007, MathWorks). Hindpaw stimulation (600 µA, 4 Hz, 1 ms pulse duration, 16 pulses) was delivered to the right hindpaw using subcutaneous needle electrodes. Hemodynamic signal was measured by imaging reflected light intensity using a CCD camera (Pixelfly, PCO) coupled to a Leica MZ16FA microscope. One trial lasted 16 s, stimulus onset was digitally triggered by a TTL pulse after a baseline recording over 4 s. Each animal was subjected to 10 stimulation trials. All these trials were averaged and signal changes were normalized to baseline.

### *Ex vivo* validation of lesion size

To validate the *in vivo* measurement of the lesion size, mice were euthanized 24 hours after the experiment by receiving an overdose i.p. injection of pentobarbital (200 mg/kg) followed by decapitation. Brains were extracted and placed in 2% 2,3,5-triphenyltetrazolium chloride (TTC, cat. #T8877, Sigma-Aldrich, St. Louis, MO) for 10 min at 37 °C to delineate infarcts which appear pale after staining. *Ex vivo* infarct areas were determined by setting a threshold of 0.5 to normalized TTC image. Specifically, pixels with intensities higher than the setting threshold were classified into infarct region. To estimate the accuracy of LMI and LSCI measurement, cross correlation coefficient between *in vivo* infarct core area and *ex vivo* infarct area was calculated for each mouse by *coefficient* = *S*_*overlapped*_/*S*_*TTC*_, where *S*_*overlapped*_ is the overlapped area between LMI/LSCI and TTC stained infarct area and *S*_*TTC*_ is the TTC-stained infarct area.

### Image processing and data analysis

Reconstruction of each LMI image was performed by simply calculating the maximum intensity projection (MIP) of the image stack including 30×30 raster scan. Quantification of blood flow information was performed based on pixel-wise fluorescence signal time course. BFI was defined as a slope of the rising fluorescence intensity signal plotted against time for 10 s after intravenous injection. The background fluorescence signal was removed resulting in the baseline signal starting from 0. The fluorescence signal profiles were normalized against the maximum intensity within each ROI time traces to compensate for any differences in actual injection dose. Linear fit was performed through the rising point and the first peak value in the time profile of each pixel, and its slope (in % s^−1^) was used to represent the BFI. TTP maps in the infarcted hemisphere were calculated as the time delay between the signal peak in each pixel with respect to the time point when the MCA became pronounced in the healthy hemisphere. The MTT maps, representing the average time of fluorescent dye perfusion in each pixel, were calculated by dividing cerebral blood volume (CBV) by BFI. CBV was calculated by the integration of fluorescence intensity over time between the wash-in and wash-out time point which were defined as the time corresponding to 10% of maximum intensity before the first peak and 40% of maximum intensity after the first peak, respectively. Quantitative flow velocity was calculated based on two points of interest in the same blood vessel as the distance between these two points divided by the latency time of their peak values in the corresponding signal time courses. Subsequently, CBV was calculated by multiplying the velocity with the respective cross-sectional area of the vessel segment. An illustration of this calculation can be found in Fig. S1. PCA was also performed to check the dynamic contrast-enhanced images. An image stack lasting 5 s post-injection including 400 frames were loaded into MATLAB software and analyzed with the embedded “princomp” function to perform PCA. The positive values from the second component were assigned a red color to represent arterial vessels. The negative values from the third component were assigned a blue color to represent veins. To better extract skull vessels, PCA was performed on the image stack within a time window between 4 – 8s and the negative values from the second components were assigned a green color.

To assess the infarct core and penumbra regions, BFI ratio between the ipsilateral and the contralateral hemispheres was calculated pixel by pixel with a sliding window of 30×30 pixels. A residual of BFI lower than 40 % was set as the threshold for the core region and 40∼60 % was set for the penumbra. Subsequently, core and penumbra regions were selected as the ROIs for functional and structural statistics. Fluorescence intensity, BFI, TTP, MTT as well as LSCI intensity were calculated as the mean pixel value inside the ROI in the corresponding functional maps and then normalized based on the contralateral side. Lesion sizes of the core and penumbra were calculated as the planar area in the corresponding regions. Vessel fill fraction or density was calculated pre- and posttreatment from the core and penumbra in the LMI time-lapse images, which was characterized as the number of pixels occupied by the cerebral vessels divided by the total number of pixels in the ROI. Before vessel density estimation, a multiscale vessel enhancement was used to differentially boost signals from small versus large vessels to remove the non-vessel background. A detailed description on vessel density estimation can be found elsewhere ^[43]^.

### Statistical analysis

Twenty mice including fourteen C57BL/6 mice and six BALB/c mice were used in this study. In each dataset, core and penumbra regions were selected based on BFI change map, followed by the computation for averaged fluorescence intensity, BFI, TTP, LSCI intensity, and vessel fill fraction in the ROIs. Data were shown in box-and-whiskers plots with median indicated as the central line inside the boxes, and interquartile range indicated as the box length. Maximum and minimum values were labeled with the whiskers. The correlation coefficients between LMI/LSCI and TTC staining passed Shapiro-Wilk normality test and were tested with paired t-test (two-tailed). A *p* < 0.05 was considered statistically significant.

## Author Contributions

ZC conceived the study; ZC, QZ, JD, CG and MA performed the *in vivo* experiments; JD performed the TTC staining; MW performed the IOI mapping; ZC, QZ and YL analyzed the data; ZC, QZ and MA interpreted the results; ZC wrote the paper; BW, SW, MA and DR supervised the project; all authors contributed constructively to the manuscript.

## Acknowledgements

The authors acknowledge grant support from the Swiss heart foundation, the Swiss National Science Foundation (SNSF) grants 310030_182703, 310030_200703, and 310030_192757, and the UZH CRPP stroke.

## Data availability

The data that support the findings of this study are available from the corresponding authors upon request.

## Code availability

The code that supports the findings of this study are available from the corresponding authors upon request.

