## Supplementary information for "Transcranial cortex-wide imaging of murine ischemic perfusion with large-field multifocal illumination fluorescence microscopy"

**Supplementary Figure 1 |** Illustration of blood flow index (BFI), time to peak (TTP), mean transient time (MTT), blood flow velocity and cerebral blood volume (CBV) calculation.

**Supplementary Figure 2 |** Comparison of cortical blood flow in the contralater hemisphere before and after stroke surgery with fluorescence localization microscopy and fluorescent beads injection.

**Supplementary Figure 3 |** LMI reveals dramatically different mean transient time (MTT) and time-to-peak (TTP) maps between different mouse strains post stroke.

**Supplementary Figure 4 |** Correlation check between BFI map and TTC image from a reprensentative mouse.

**Supplementary Figure 5 |** Correlation check between MTT map and TTC image from a reprensentative mouse.

**Supplementary Figure 6 |** Correlation check between TTP map and TTC image from a reprensentative mouse.

**Supplementary Figure 7 |** Immunohistochemistry of an infarct brain slice with NeuN and DAPI staining.

**Supplementary Figure 8 |** MRI data coregistration between T1 weighted image and MR angiogram and fMRI BOLD images based on SPM12.

**Supplementary Figure 9 |** Mouse brain perfusion validation with MRI and superparamagnetic iron dioxide nanoparticles.

**Supplementary Video 1 |** Perfusion dynamics from a representative BALB/c mouse post stroke.

**Supplementary Video 2 |** Perfusion dynamics from a representative C57BL/6 mouse post stroke.

**Supplementary Note 1 |** Cerebral blood flow imaging with widefield fluorescence localization microscopy.

**Supplementary Note 2 |** Cerebral blood flow imaging with dynamic contrast-enhanced perfusion magnetic resonance imaging using superparamagnetic iron dioxide nanoparticles (SPION)

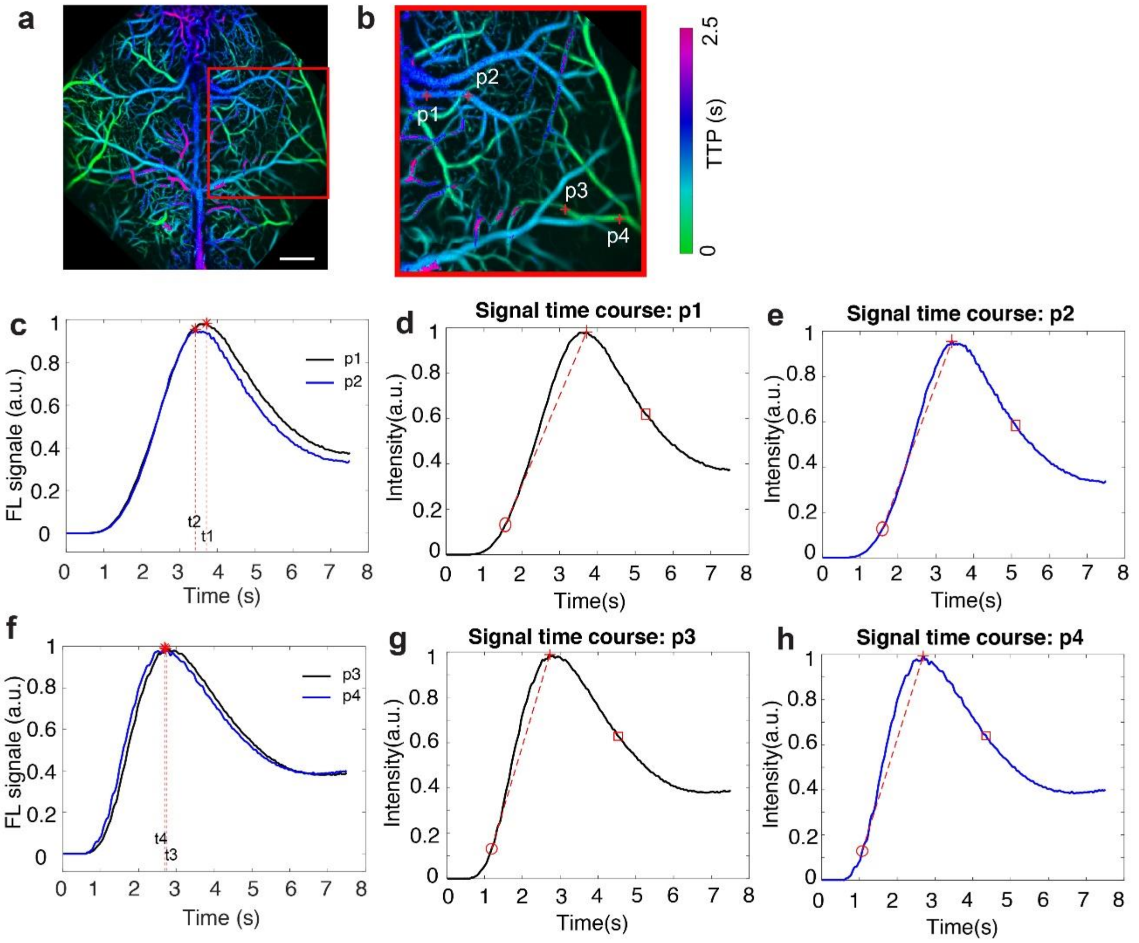


**Supplementary Figure 1 |** Illustration of blood flow index (BFI), time to peak (TTP), mean transient time (MTT), blood flow velocity and cerebral blood volume (CBV) calculation. (a) TTP map from a representative mouse. (b) Zoom-in view of the red boxed region in (a). p1 – p4: points of interest selected to check BFI, TTP, MTT, flow velocity and CBV. (c) Signal time course from p1 and p2. Red dotted lines indicate the latency time between the signal peaks. **$V_{p1\text{-}p2}$ = distance$_{p1\text{-}p2}$ / (t2 – t1)** = -2.0614 mm/s, indicating the flow direction is from p2 to p1 with a velocity of ~2.06 mm/s. Correspondingly, **CBV = $V_{p1\text{-}p2}$ * π *diameter^2/4** = -0.011703 mm^3/s. (d), (e) Signal time courses from p1 and p2, respectively. The red circle indicates the starting time point for BFI and MTT calculation, when the signal reaches 10% of its peak value. The red cross indicates the peak time point; the red squre indicates the end time point for MTT calucation when the signal drops by 60% from the peak to post-injection signal level after 10s. BFI is calcuated as the slope of linear signal increase from starting time and the peak, i.e., $BFI = \frac{sig(t_{peak})-sig(t_{start})}{t_{peak}-t_{start}}$; TTP is calcuated as the latency time between the peak and the starting time point from the middle cerebral artery, i.e, $TTP = t_{peak} - t_{MCA\_start}$; MTT is calculated as $MTT = \sum_{t\_start}^{t\_end} \frac{\mathrm{sig(t)}}{\mathrm{frame\_rate}} / \mathrm{BFI}$. (f) - (h) Corresponding signal time courses of p3 and p4. $V_{p3\text{-}p4}$ = -23.1777mm/s and CBV = -0.042966 mm^3/s.

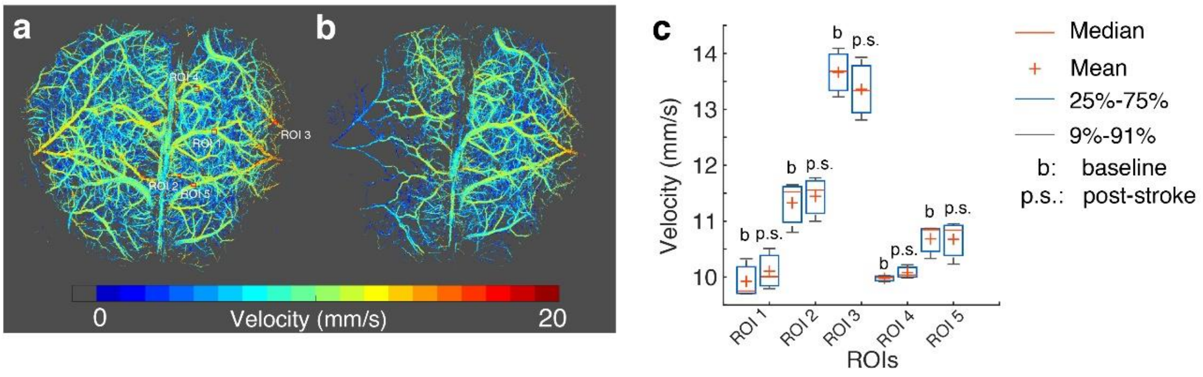


**Supplementary Figure 2 |** Comparison of cortical blood flow in the contralateral hemisphere before and after stroke surgery. (a) Cerebral blood flow velocity map before stroke (baseline) acquired with a fluorescent localization microscopy system with fluorescent beads injection. For more details, please check **Supplementary Note 1**. (b) Corresponding blood flow velocity map after stroke. (c) Velocity comparison from 5 regions of interests (ROIs) as shown in panel (a). No significant velocity changes were observed from the contralateral side before and after stroke.

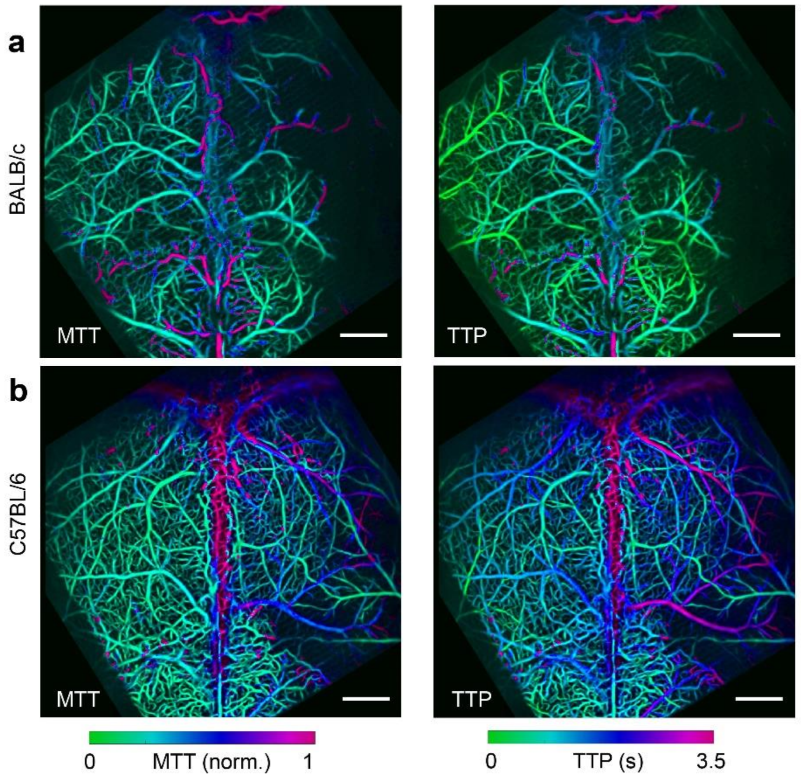


**Supplementary Figure 3 |** LMI reveals dramatically different mean transient time (MTT) and time-to-peak (TTP) maps between different mouse strains post stroke. **a** MTT and TTP map from a representative BALB/c mouse. **b** Corresponding results from a representative C57BL/6 mouse. All scale bars: 1 mm.

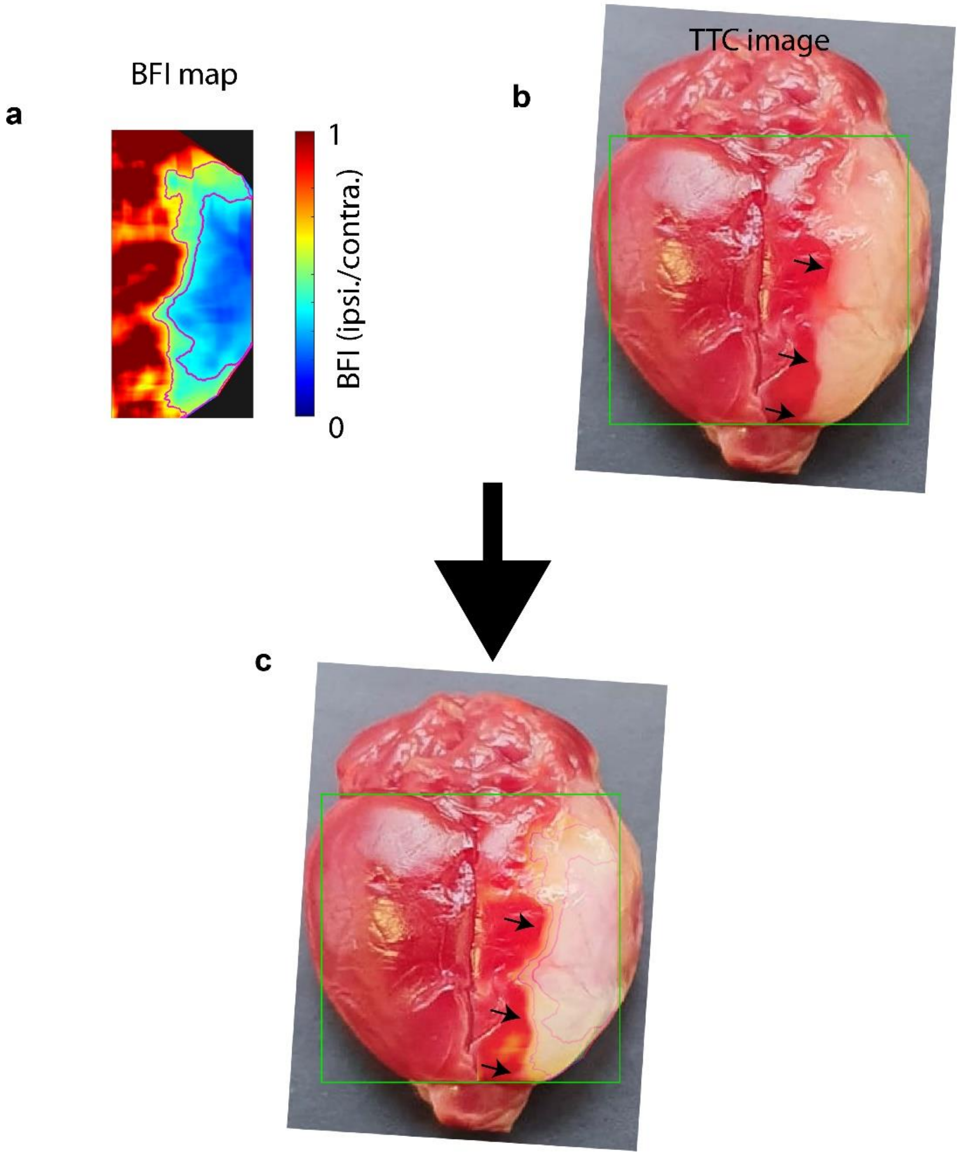


**Supplementary Figure 4 |** Correlation check between BFI map and TTC image from a reprensentative mouse. (a) BFI map. (b) TTC image. (c) BFI overlaid on the TTC image. The bounday of penumbra shown in BFI map nicely matches the lesion area in the TTC image. BFI map can distinguish the infarct core and peneumbra when a residual of BFI lower than 40 % is set as the threshold for the core region and 40~60 % is set for the penumbra.

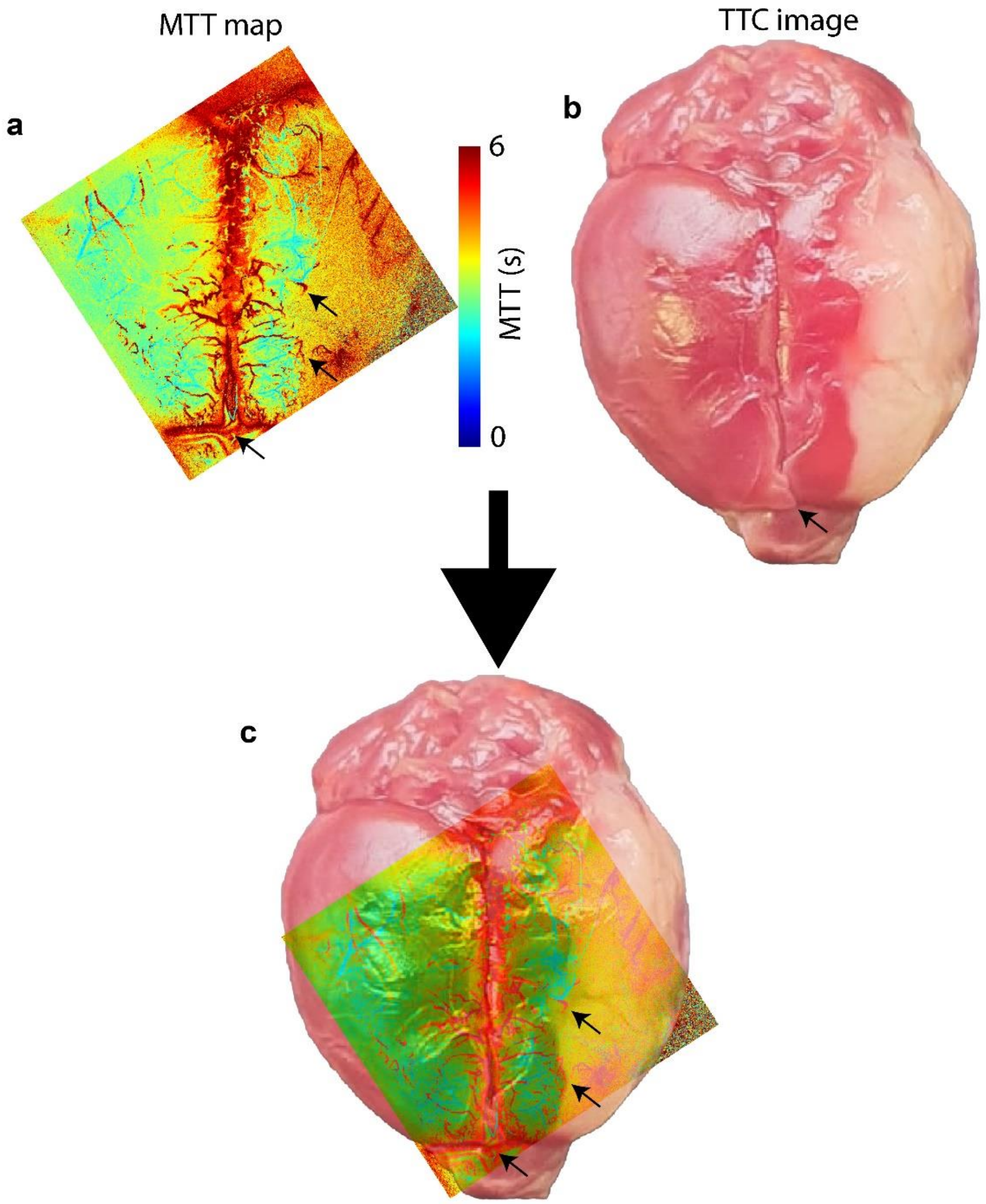


**Supplementary Figure 5 |** Correlation check between MTT map and TTC image from a reprensentative mouse. MTT map presents clear boundary between health and infarct tissue as infarct tissue has prolonged MTT values due to impaired microcirculation. However, MTT map is inferior to distinguish the infarct core and penumbra.

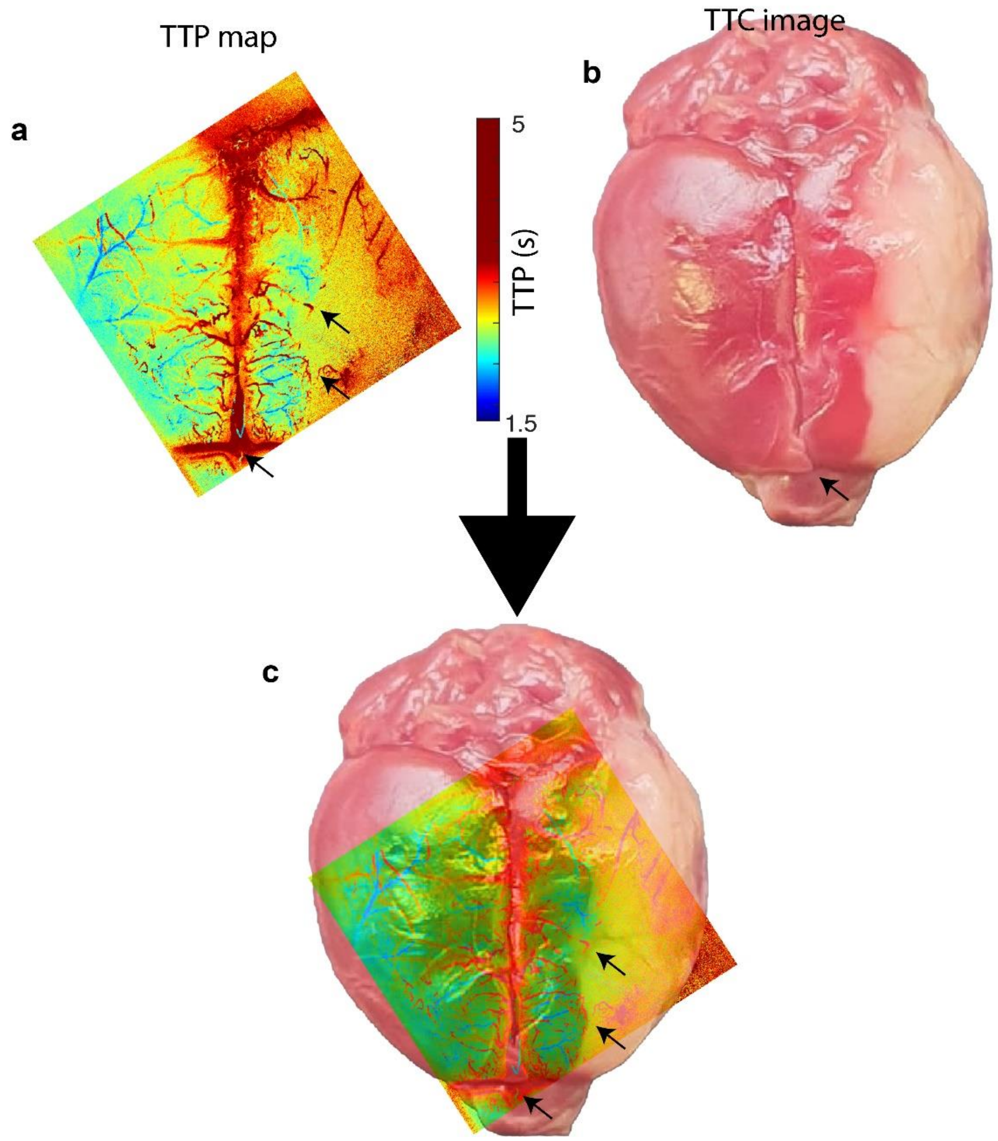


**Supplementary Figure 6 |** Correlation check between TTP map and TTC image from a reprensentative mouse. Similar to MTT map, TTP map presents clear boundary between health and infarct tissue as infarct tissue has prolonged TTP values due to impaired microcirculation. However, same as MTT map, TTP map is inferior to distinguish the infarct core and penumbra.

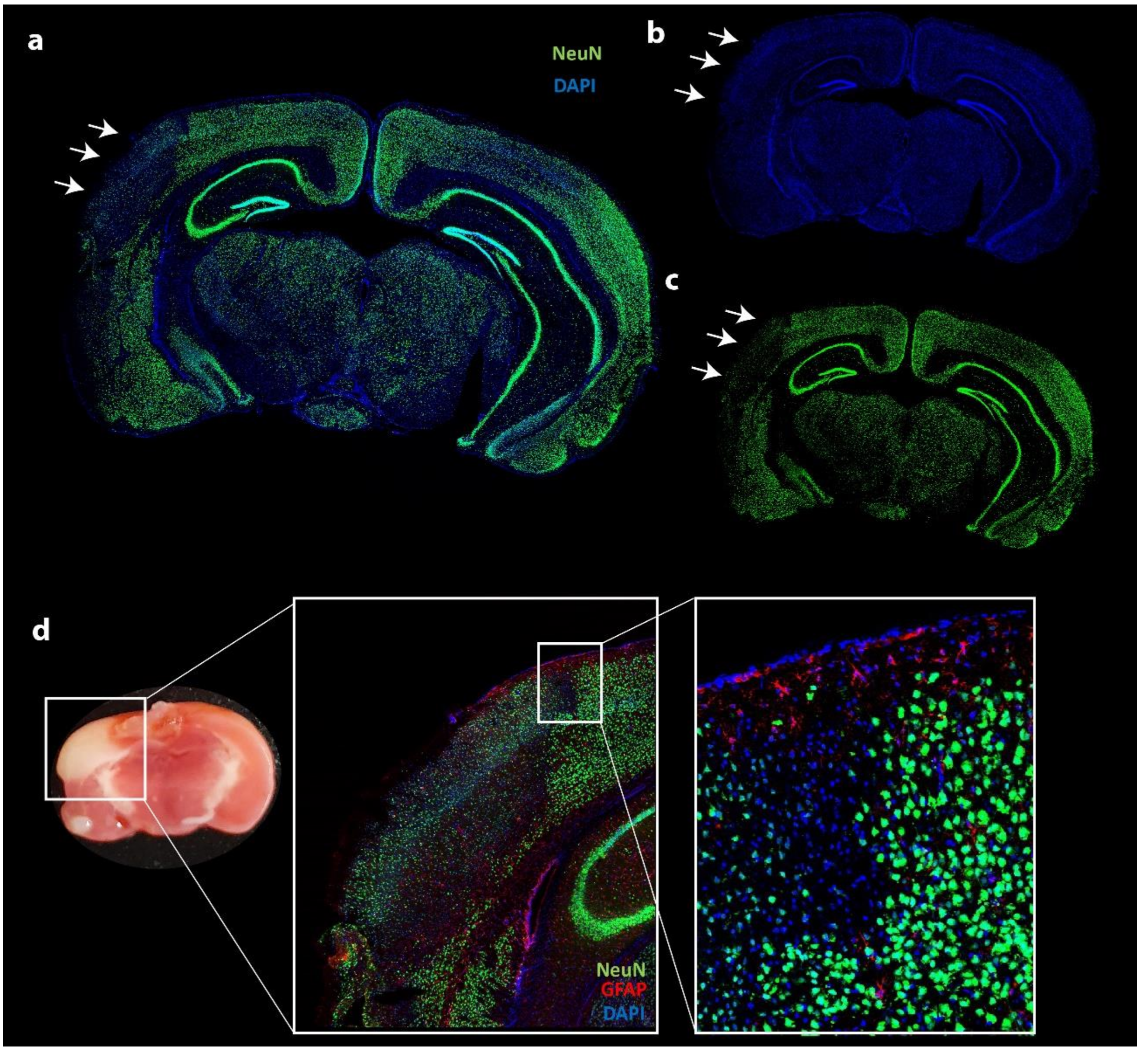


**Supplementary Figure 7 |** Immunohistochemistry of an infarct brain slice with NeuN and DAPI staining. (a) Superimposed image of NeuN and DAPI channels. White arrows indicate the infarct region post stroke. (b) DAPI staining (general cell marker) doesn't show obvious different between two hemisperes. (c) NeuN staining (neuro-specific marker) clearly shows reduced neuron density in the infarct region. (d) TTC staining (tissue damage marker) correlates well with immunohistochemistry of the infarct region dispite of some tissue morphological changes during the staining processes.

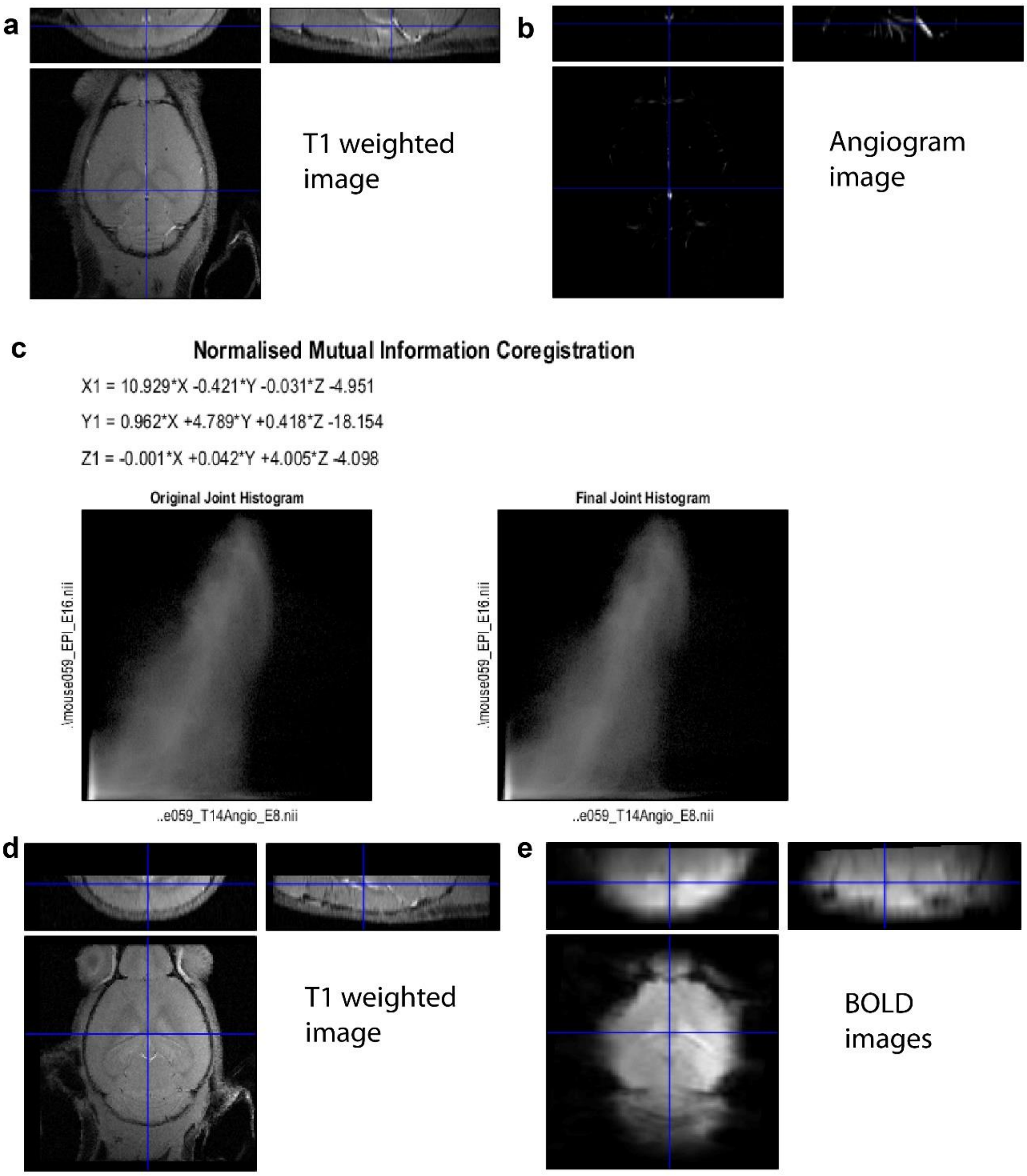


**Supplementary Figure 8 |** MRI data coregistration between T1 weighted image and MR angiogram and fMRI BOLD images based on SPM12 [2].

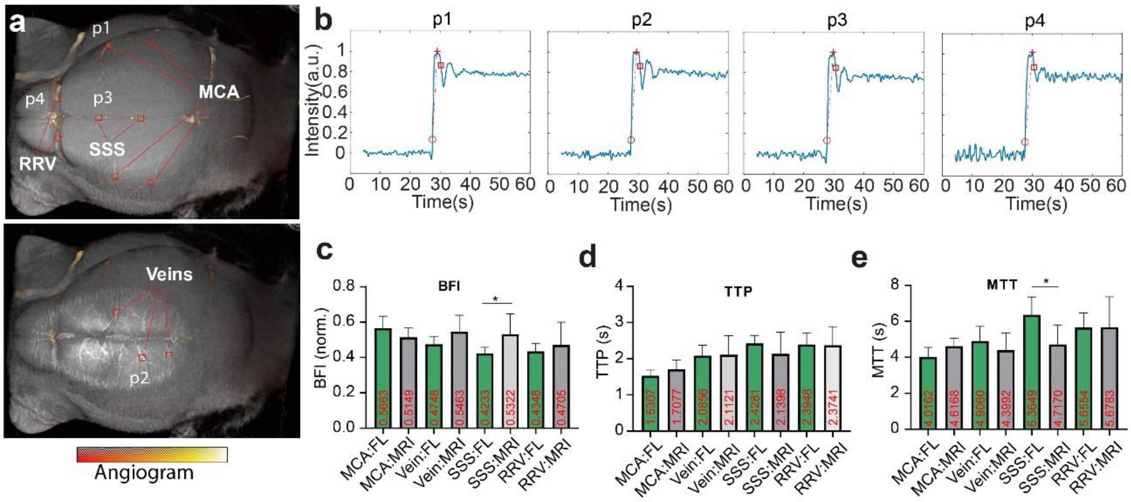

**Supplementary Figure 9 |** Mouse brain perfusion validation with MRI and superparamagnetic iron dioxide nanoparticles (SPIONs). (a) Anatomical image of the mouse brain. MR angiogram acquired with 2D FLASH sequence was overlaid on the T1 weighted image. p1 – p4: points of interest to check perfusion parameters. For more details, please check **Supplementary Note 2.** (b) Signal time course from p1-p4 from blood oxygen level dependent (BOLD) signals acquired with the EPI sequence. The signal was inverted by subtracting the signal post injection from the baseline. BFI, TTP and MTT calculation was same as the fluoresence data. $BFI_{p1}$ = 0.5547, $TTP_{p1}$ = 1.5588 s, $MTT_{p1}$ = 4.4078; $BFI_{p2}$ = 0.4666, $TTP_{p2}$ = 2.1774 s, $MTT_{p2}$ = 5.8276; $BFI_{p3}$ = 0.4185, $TTP_{p3}$ = 2.5348 s, $MTT_{p3}$ = 6.1172; $BFI_{p4}$ = 0.3626, $TTP_{p4}$ = 2.7295 s, $MTT_{p4}$ = 7.1919. (c)-(e) BFI/TTP/MTT comparison between fluorescence (n = 3 mice) and MRI measurements (n = 3 mice) based on ROIs selected in different vessels as shown in (a). The difference is not significant in each group except BFI and MTT from SSS, where fluorescence data show lower BFI and longer MTT arguably caused by fluorescence signal intengration across different depths such as cortex and skull. Statistics was done with non-parametric Mann-Whitney test, and $p<0.05$ was considered statistically significant. MCA: middle cerebral artery; SSS: superior sagittal sinus; RRV: rostral rhinal vein.

### Supplementary Note 1

Cerebral blood flow imaging before and after stroke surgery was performed with a previously developed widefield fluorescence localization microscopy (WFLM) system [1].

##### Experimental set-up of the WFLM system

The WFLM system was comprised of a continuous wave (CW) laser sources at 473 nm (FPYL-473-1000-LED, Frankfurt Laser Company, Germany) for fluorescence excitation and a high-speed CMOS camera for fluorescence detection. The laser beam was coupled into a commercial fiber bundle to provide epi-illumination. The backscattered fluorescence was collected with a commercial macroscopic objective (AF micro-Nikkor 105 mm, Nikon, Japan) and filtered with a long-pass filter (FGL515, Thorlabs, USA) before entering the sensor plane of the CMOS camera (pco.dimax S1, PCO AG, Germany). The high-speed camera features a fast frame rate up to 4.4 kHz at full pixel resolution of 1008 × 1008 pixels. During data acquisition, a frame rate of 400 Hz was selected after taking multiple factors into consideration, such as image signal to noise ratio, measured blood flow velocity range and injected fluorescent microspheres. Image acquisition was performed with Camware software (version 4.12, PCO AG, Germany).

##### *In vivo* animal experiment

In vivo animal experiment was performed to a 3-month old C57BL/6 mouse. Before and after stroke surgery, cerebral blood flow was mapped with WFLM by injecting the orange-yellow fluorescent beads (FMOY-1.3, 460/594 nm, Cospheric, U.S.) with 1–5 µm diameter diluted in water administered intravenously through the mouse tail vein. The mouse was anesthetized with isoflurane (3.0% v/v for induction and 1.5% v/v during experiments) in 20%O2 and 80% air at a flow rate of ~1l/min). During the experiment, the skull of the mouse was kept intact, while the scalp was removed to reduce light scattering. Mouse physiological status was monitored in real-time with Physiosuite (Kent Corp., USA) and the body temperature was kept around 37 °C.

##### Image reconstruction and velocity map calculation

The principle of image processing is based on localization microscopy and particle tracking algorithms. The fluorescent emitters in each frame were localized by searching for local maxima with an adaptive threshold method, while the centroid coordinates of blurring point spread functions (PSFs) were recorded corresponding to different fluorescent bead locations. Subsequently, the trajectory of each recognizable emitter was tracked with a Simpletracker algorithm. Two main parameters are involved to establish the linking of the same emitter in time-lapse frames: max-linking-distance (MLD) and max-gap-closing (MGC). The MLD parameter defines the acceptable displacement to search for the same beads between consecutive frames, which is dependent on the frame rate and potential flow velocity range. An empirical threshold of linking length was used to preserve only effective linking. The localized image was subsequently rendered by superimposing all the particle trajectories, while the brain-wide maps of the flow velocity and direction were reconstructed by considering the relative displacement of each bead for the given fixed frame rate.

#### Supplementary Note 2

Cerebral blood flow measurement with dynamic contrast-enhanced perfusion magnetic resonance imaging using superparamagnetic iron dioxide nanoparticles (SPION)

##### Experimental set-up

The dynamic contrast-enhanced perfusion magnetic resonance imaging (MRI) experiment was conducted on a 7 T Bruker Biospec 70/16 small animal MR system (Bruker BioSpin MRI, Ettlingen, Germany) using a T/R Cryocoil. ParaVision 6.0.1 was used as the user interface.

##### ***In vivo*** animal experiment

Three 8-week old female C57BL/6 mice were imaged non-invasively to validate the measured perfusion parameters from the proposed LMI method. Each mouse was anesthetized with isoflurane (3.0% v/v for induction and 1.5% v/v during experiments) in 20%O2 and 80% air at a flow rate of ~1l/min). Mouse body temperature was kept around 37 °C using a water heating system throughout the experiment. Superparamagnetic iron dioxide nanoparticles (SPION, Endorem, AMAG Pharmaceuticals, Inc., concentration 11.2mg Fe/ml) were adminstratered manually and intravenously through the mouse tail vein (40 µl injection volume). Functional MRI data was recorded 100 s before contrast agent injection.

##### MRI data acqusition and data analysis

MRA images were acquired using a FLASH sequence: FOV = 16 × 16 mm$^2$, matrix dimension (MD) = 320 × 320, 20 slices from the brain surface to deeper regions, slice thickness = 0.3 mm, repetition time (TR) = 7.3 ms, echo time (TE) = 2.0 ms, number of averages (NA) = 4. T1-weighted scan in the transverse plane was acquired as anatomical reference for the fMRI data using a FLASH sequence: FOV = 16 × 16 mm$^2$, MD = 384 × 384, 20 slices from anterior to posterior, slice thickness = 0.3 mm, TR = 287.1 ms, TE = 3.5 ms, NA = 3. Prior to fMRI data acquisition, the local field homogeneity was optimized using the acquired B0 field maps. BOLD data were acquired using gradient-echo echo-planar imaging (GE-EPI) sequence: FOV = 18

× 18 $mm^2$, MD = 70 × 70, yielding an in-plane voxel dimension of 257 × 257 $\mu m^2$, 10 slices from surface to bottom, slice thickness = 0.2 mm, flip angle (FA) = 60°, TR = 400 ms, TE = 0.28 ms, NA = 1, yielding an effective temporal resolution of 2 Hz for the volumetric acquisitions.

MRI data from each mouse were firstly coregistered between T1 weighted image, MR angiogram and fMRI BOLD images based on SPM12 software. Subsequently, blood perfusion dynamics were characterized by calculating the BFI, MTT and TTP from the middle cerebral artery, large drainage veins, superior sagittal sinus and the rostral rhinal vein in the same way as done to the fluorescence data. BFI/TTP/MTT comparison between fluorescence (n = 3 mice) and MRI measurements (n = 3 mice) based on the above mentioned ROIs was plotted and statistically tested with non-parametric Mann-Whitney test.

[2] The Wellcome Centre for Human Neuroimaging https://www.fil.ion.ucl.ac.uk/spm/software/spm12/
